# Allelic gene polymorphisms suspected to diversify the individual early metabolic response upon influenza H3N2 and SARS-CoV-2 infections

**DOI:** 10.1101/2022.05.05.490776

**Authors:** Birgit Arnholdt-Schmitt, Shahid Aziz, José Hélio Costa

**Affiliations:** Non-Institutional Competence Focus (NICFocus) ‘Functional Cell Reprogramming and Organism Plasticity’ (FunCROP), coordinated from Foros de Vale de Figueira, Alentejo, Portugal; Functional Genomics and Bioinformatics, Department of Biochemistry and Molecular Biology, Federal University of Ceara, 60451-970, Fortaleza, Ceara, Brazil

## Abstract

In this brief report, we point to virus infection time-dependent transcript levels of polymorphic ASMTL genes in human nasal epithelial cells from seven cell origins. Our observations encourage focused top-down and hypothesis-driven studies in support of more efficient allelic genotyping to identify targets for early resilience prediction.

---

Recently, we identified early ROS/RNS rebalancing as part of a major complex trait, named CoV-MAC-TED that characterized host cell response upon SARS-CoV-2 infections from two genetic virus variants (1, 2, 3). Transcript level changes of the enzymes acetylserotonin O-methyltransferase-like (ASMTL) and alcohol dehydrogenase 5 (ADH5) were hypothesized to mark relevant adaptive ROS/RNS equilibration early during respiratory infections and preliminary validation in cultured primary target cells supported this view (2). ASMTL is a paralog of ASMT, which is involved in final melatonin biosynthesis. ADH5 also known as S-nitrosoglutathione reductase (GSNOR) is involved in NO homeostasis (references cited in 1 and 2). In parallel to these studies, we released our results on *ASMTL* and *ADH5* transcriptome accumulation from influenza H3N2-infected human nasal epithelial cells (NECs) that originated from seven healthy cell donator origins (D01-D07) (4). We observed that variability of *ASMTL* and *ADH5* transcript levels linked to the timing of virus replication and that this connected to the initiation of the classical immune system response, when marked by interferon regulatory factor 9 (*IRF9*) transcript level changes. Especially, cells of the donator D02 showed at 8 hours post infection (hpi) highest *ASMTL* transcript levels among all cell origins indicating rapidly unbalanced ROS/RNS, which linked to the earliest immune response. On the contrary, transcript levels of *ADH5* from D05 indicated early and over times highest nitric oxide (NO) stress among all seven origins, suggesting unbalanced ROS/RNS in favor of RNS. This response connected to the highest immune response at 24 hpi among all cell origins. In contrast, cells originating from D01 indicated a stably balanced relationship between ROS and RNS marked by *ASMTL* and *ADH5* that associated to the lowest *IRF9* response over all time points among the seven cell origins. This equilibrated response of D01 was in accordance with postponed influenza virus replication in relation to all other cell origins (4). Of note, also under infection by two SARS-CoV-2 variants NECs from donator origin D01 displayed contrasting response to other donator cells: lower transcript levels for enolase (representing glycolysis) and lactate dehydrogenase (LDH) (representing aerobic fermentation) were observed at 24 hpi for cell cultures from origin D01 that linked to delayed replication of SARS-CoV-2 (3).

These overall observations made us optimistic that our approach and marker systems in combination with appropriate primary target human cells could be promising, in general, to early identify differential individual host cell responses upon viral infections. At the same time, we hypothesized that this tool could also help to discriminate individuals through genetic polymorphisms in the relevant functional genes that relate to CoV-MAC-TED (see ReprogVirus gene set in 1, 2, 3, and 4).

ROS signaling is one of the earliest and relevant cell responses upon abiotic and biotic stresses that transmits any change in environment, including viral attacks, to structures and molecules (membranes, ion channels (Ca^+^), enzymes, microtubulines etc), which then are crucially determining cell performance. Therefore, we studied polymorphisms in both highlighted marker genes for ROS/RNS balancing, *ASMTL* and *ADH5.* In a first step, we checked the genetic material of the public transcriptome data of these seven donators for NECs (5). We realized with surprise that the coding sequence of *ADH5* was highly conservative. No sequence nucleotide polymorphism (SNP) or other major differences among the seven donator cell origins were encountered. To the contrary, we found several polymorphic sites (SNPs) in the coding sequence of the *ASMTL* gene (unpublished data). This might indicate a higher potential for acclimation acquired during evolution.

In this brief report, we want to point to transcript level profiles that we observed for one of the polymorphic sites during early virus-induced cell reprogramming as an example to highlight the general importance of this finding. In the upper part of **table 1**, we demonstrate the crucial nucleotide for polymorphism in this site (G or A) for influenza H3N2-infected cells from seven donator origins. On the one hand, we observed cell origins with the same preferential nucleotide profile over times in this site: e.g. cells of D02, D03 and D07 show continuously G at all time points (0, 8 and 24 hpi); D04 and D06 display stably A, such as in the reference gene. On the other hand, we discovered for D01 and D05 unique differential transcript level profiles. Cells from D05 changed at 8 hpi to the preferential transcription of an allel, which shows A instead of G in this polymorphic site. However, at 24 hpi a change in preferential allelic transcription became visible through the reappearance of G instead of A. In contrast, at 8 hpi D01 changed in the same way as observed for D05 to preferential transcription of an allel that shows A instead of the initial G. However, in cells of D01 this allelic site continued longer to be preferentially transcribed and A is still the more frequent nucleotide seen at 24 hpi (dark green at 8 hpi and 24 hpi). Post infection time-dependent allelic genotyping profiles of the same polymorphic site was also found upon infection by two SARS-CoV-2 variants that demonstrated diverse effects on disease severity (3). ASMTL genotype profiles are shown for NECs from D01 and D03 (table 1, lower part). Overall, these results demonstrate virus- and variant-specific allelic genotype profiles over times when both virus types (Influenza H3N2 and SARS-CoV-2) and both SARS-CoV-2 variants are compared within each cell origin.

**Table 1:**
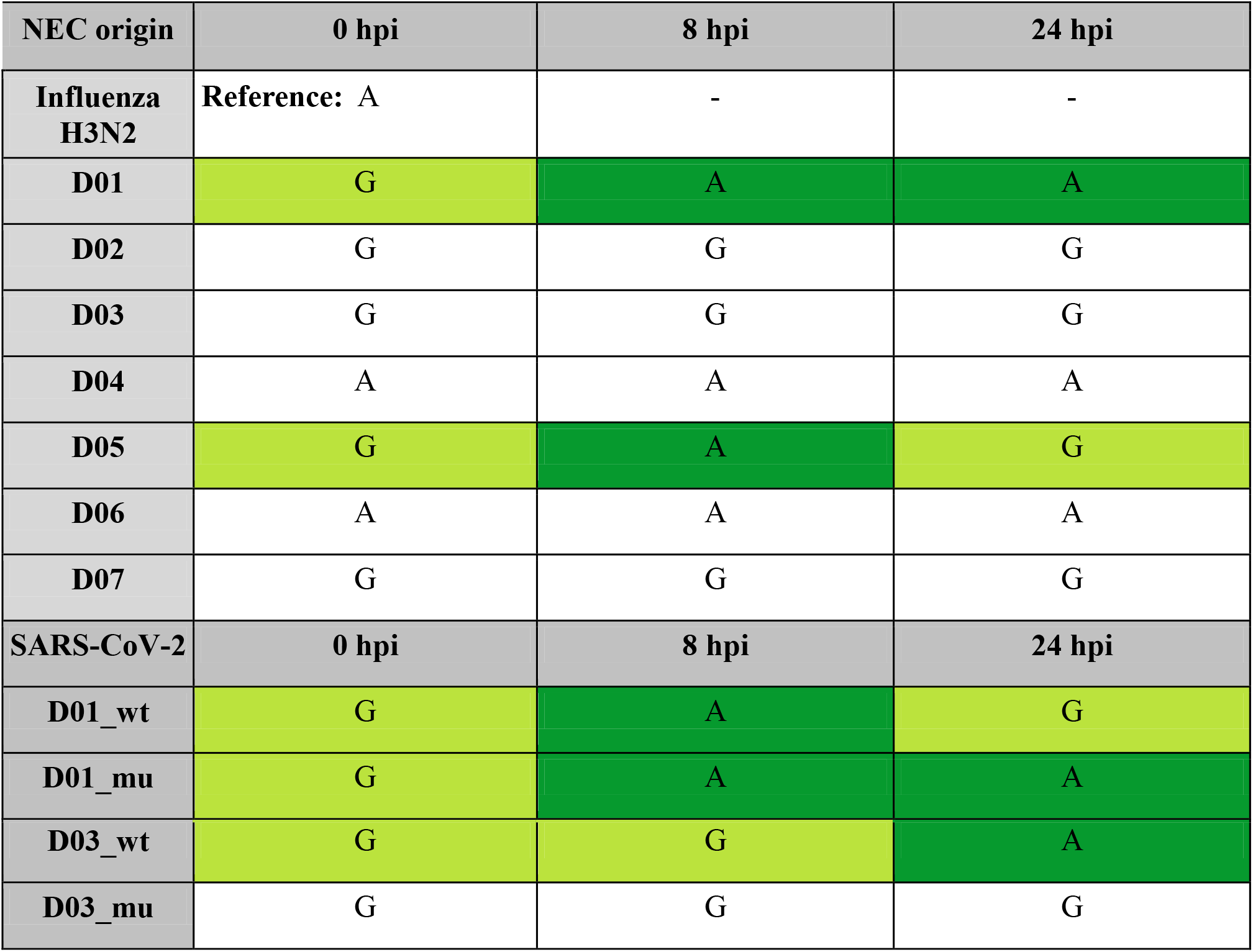
NEC origin-dependent differential transcript levels of polymorphic ASMTL genes at 8 hpi and 24 hpi upon infection with influenza H3N2 and two SARS-CoV-2 variants. (unpublished data) Wt: wild type, refers to originally identified SARS-CoV-2, mu: refers to a mutated SARS-CoV-2 (3); changes in preferential transcription of allels that demonstrate G or A in the polymorphic position at 0 hpi, 8 hpi and 24 hpi are highlighted through a switch in the table cell color from light to dark green and vice versa.

| NEC origin | 0 hpi | 8 hpi | 24 hpi |
| --- | --- | --- | --- |
| **Influenza H3N2** | Reference: A | - | - |
| **D01** | G | A | A |
| **D02** | G | G | G |
| **D03** | G | G | G |
| **D04** | A | A | A |
| **D05** | G | A | G |
| **D06** | A | A | A |
| **D07** | G | G | G |
| **SARS-CoV-2** | **0 hpi** | **8 hpi** | **24 hpi** |
| **D01_wt** | G | A | G |
| **D01_mu** | G | A | A |
| D03_wt | G | G | A |
| D03_mu | G | G | G |

In summary and despite the restricted number of studied indivudual cell origins, these observations highlight to our view that efficient allelic genotyping for specific or general virus tolerance and immunophenotyping requires integrative studies on time-dependent individual transcript level changes in genes that were identified as relevant markers for early cell reprogramming, such as recently reported for CoV-MAC-TED. Rapid switching between critical gene polymorphisms in critical binding sites can determine metabolic regulation, which in turn can result in differential performance. This was described in plant research for the ROS/RNS balancing gene alternative oxidase (*AOX*) and its temperature-dependent interaction with pyruvate (6). Furthermore, we suggest that high diversity in coding and non-coding sequences of crucial genes for early ROS/RNS balancing should be explored as a trait *per se.* This trait can support plasticity in the immune response. Consequently, we propose a discussion on a potential paradigm shift towards understanding immunology in a wider sense, which should consider ROS/RNS balancing during stress-induced early cell reprogramming. It is our hope that our rapid communication of these insights might help to avoid spending high amounts of personal and financial resources for less-focused bottom-up gene polymorphism studies instead of top-down and then hypothesis-driven approaches (7, 8).

## References

1. Arnholdt-Schmitt B, Mohanapriya G, Bharadwaj R, Noceda C, Macedo ES, Sathishkumar R, Gupta KJ, Sircar D, Kumar SR, Srivastava S, Adholeya A, Thiers KL, Aziz S, Velada I, Oliveira M, Quaresma P, Achra A, Gupta N, Kumar A, Costa JH. From Plant Survival Under Severe Stress to Anti-Viral Human Defense - A Perspective That Calls for Common Efforts. Front Immunol. 2021 Jun 15;12:673723. doi: 10.3389/fimmu.2021.673723.

2. Costa JH, Mohanapriya G, Bharadwaj R, Noceda C, Thiers KLL, Aziz S, Srivastava S, Oliveira M, Gupta KJ, Kumari A, Sircar D, Kumar SR, Achra A, Sathishkumar R, Adholeya A, Arnholdt-Schmitt B. ROS/RNS Balancing, Aerobic Fermentation Regulation and Cell Cycle Control - a Complex Early Trait (‘CoV-MAC-TED’) for Combating SARS-CoV-2-Induced Cell Reprogramming. Front Immunol. 2021 Jul 7;12:673692. doi: 10.3389/fimmu.2021.673692

3. Costa JH, Aziz S, Noceda C, Arnholdt-Schmitt B. Major Complex Trait for Early De Novo Programming ‘CoV-MAC-TED’ Detected in Human Nasal Epithelial Cells Infected by Two SARS-CoV-2 Variants Is Promising to Help in Designing Therapeutic Strategies. Vaccines (Basel). 2021 Nov 26;9(12):1399. doi: 10.3390/vaccines9121399

4. Costa JH, Aziz S, Noceda C, Arnholdt-Schmitt B. Transcriptome data from human nasal epithelial cells infected by H3N2 influenza virus indicate early unbalanced ROS/RNS levels, temporarily increased aerobic fermentation linked to enhanced α-tubulin and rapid energy-dependent IRF9-marked immunization. https://www.biorxiv.org/content/10.1101/2021.10.18.464828v2. Posted November 09, 2021

5. Gamage AM, Tan KS, Chan WOY, Liu J, Tan CW, Ong YK, Thong M, Andiappan AK, Anderson DE, Wang Y, Wang LF. Infection of human Nasal Epithelial Cells with SARS-CoV-2 and a 382-nt deletion isolate lacking ORF8 reveals similar viral kinetics and host transcriptional profiles. PLoS Pathog. 2020 Dec 7;16(12):e1009130. doi: 10.1371/journal.ppat.1009130.

6. Ito K, Ogata T, Seito T, Umekawa Y, Kakizaki Y, Osada H, et al. Degradation of Mitochondrial Alternative Oxidase in the Appendices of Arum Maculatum. Biochem J (2020) 477(17):3417–31. doi: 10.1042/BCJ20200515

7. Arnholdt-Schmitt B, Costa JH, de Melo DF. AOX - a functional marker for efficient cell reprogramming under stress? Trends Plant Sci. 2006 Jun;11(6):281–7. doi: 10.1016/j.tplants.2006.05.001.

8. Crespo I, Götz L, Liechti R, Coukos G, Doucey MA, Xenarios I. Identifying biological mechanisms for favorable cancer prognosis using non-hypothesis-driven iterative survival analysis. NPJ systems biology and applications. 2016 Dec 22;2(1):1–1. doi: 10.1038/npjsba.2016.37. eCollection 2016

